# Robust Bayesian analysis of animal networks subject to biases in sampling intensity and censoring

**DOI:** 10.1101/2024.10.01.616020

**Authors:** Sebastian Sosa, Mary Brooke McElreath, Daniel Redhead, Cody T. Ross

## Abstract

Data collection biases are a persistent issue for studies of social networks. This issue has been particularly important in Animal Social Network Analysis (ASNA), where data are unevenly sampled and such biases may potentially lead to incorrect inferences about animal social behavior. Here, we address the issue by developing a Bayesian generative model, which not only estimates network structure, but also explicitly accounts for sampling and censoring biases. Using a set of simulation experiments designed to reflect various sampling and observational biases encountered in real-world scenarios, we systematically validate our model and evaluate it’s performance relative to other common ASNA methodologies. By accounting for differences in node-level censoring (i.e., the probability of missing an individual interaction.), our model permits the recovery of true latent social connections, even under a wide range of conditions where some key individuals are intermittently unobserved. Our model outperformed all other existing approaches and accurately captured network structure, as well as individual-level and dyad-level effects. Antithetically, permutation-based and simple linear regression aprroaches performed the worst across many conditions. These results highlight the advantages of generative network models for ASNA, as they offer greater flexibility, robustness, and adaptability to real-world data complexities. Our findings underscore the importance of generative models that jointly estimate network structure and adjust for measurement biases typical in empirical studies of animal social behaviour.

## Introduction

Over the past 50 years, network analysis has become a vital tool for studying behavior in natural and artificial systems, in fields as diverse as sociology (Carrington et al. 2005; Scott 2002; Snijders 2011), anthropology (Jang et al. 2024; Power 2017; von Rueden et al. 2019), economics (Jackson 2010; Ter Wal and Boschma 2009), and ecology (Pinter-Wollman et al. 2014; Sosa et al. 2021a). Network analysis methods can be broadly applied to data at all scales—from proteomics (Ravasz et al. 2002), to animal social relationships (Croft et al. 2008), to ecosystems as a whole (Ulanowicz et al. 2014). However, in each domain, researchers must be careful to adapt general-purpose methods to deal with specific measurement issues known to affect their study systems. For example, using human social network data, social scientists routinely find that the two members of a dyad disagree on a given resource transfer—for example: individual *i* may report giving money to individual *j*, but individual *j* might not report receiving money from individual *i*. In order to make sense of such data, complex measurement models (e.g., Butts 2003; Young et al. 2020; Redhead et al. 2023b; De Bacco et al. 2023) that estimate and account for reporting error need to be explicitly linked to—that is, applied simultaneously with—standard network analysis models (Kenny and La Voie 1984; Back and Kenny 2010; Karrer and Newman 2011) that assume internally coherent raw data.

In the realm of animal sociality research, researchers have identified similar types of threats to validity, which are often caused by measurement difficulties—including sampling biases (i.e. heterogeneity and bias in sampling intensity across individuals), and censoring baises (i.e. detectability or censoring issues, where key network ties may be cryptic or concealed). In an attempt to deal with these methodological challenges, researchers have typically relied on a two step process. In the first step, dyadic sociality indices (Whitehead 1999) are constructed on the basis of focal follow or scan sampling data. In the second step, these sociality indices are included in models that use network permutation methods (Farine 2013; Farine and Whitehead 2015a), such as the quadratic assignment procedure (i.e., QAP; Hubert and Schultz 1976; Krackardt 1987) or its extensions (e.g., MRQAP; Dekker et al. 2003), to ‘account for’ the non-independence of observations. Although this two-step process was once thought to appropriately address measurement challenges in the field, recent studies have revealed significant reliability concerns associated with these hypothesis testing protocols common to studies of animal social networks (Weiss et al. 2021; Puga-Gonzalez et al. 2021; Hart et al. 2022; Farine and Carter 2022).

Such critiques have been both empirically motivated— e.g., by applying standard analysis pipelines to simulated data and finding that routinely used tools cannot recover the true parameter values used to simulate data (Puga-Gonzalez et al. 2021)—and motivated by purely mathematical considerations—e.g., standard ASNA pipelines applied to sociality indices violate the laws of probability theory (Jaynes 2003), and thus are erroneous and ineffectual *in principle*, as we will show in detail below. Similarly, the permutation methods commonly used to account for interdependence in animal social networks were sharply critiqued by one of their founders more than three decades ago as: “inappropriate on logical grounds”, “difficult to interpret”, and “biased”, with the further caveat that “…the bias is not [even] consistent but rather can be liberal or conservative, small or large, depending on parameters in the population from which the data are sampled” (Krackhardt 1992, p. 1). Although some improvements to permutation methods have been made since (Dekker et al. 2003; Hunter et al. 2008), the general shortcomings of the method remain.

As far back as the early 1990s, network scientists and applied statisticians were well aware that permutation methods do not account for interdependence in social networks— animal or otherwise—and so a suite of highly parameterized correlated random effects models—e.g., the social relations model (Kenny and La Voie 1984; Snijders and Kenny 1999; Back and Kenny 2010), stochastic blockmodels (Pearl and Schulman 1983; Holland et al. 1983; Karrer and Newman 2011; Peixoto 2019), and additive and multiplicative effects network models (Hoff 2021)—were developed to generatively model network relations. These generative network models, however, remained under-utilized for decades, largely because estimation of their parameters was computationally infeasible until only recently, with the advent of efficient Markov Chain Monte Carlo samplers, like Stan (Stan Development Team 2021). However, in the last five years, a number of open-source software packages for implementing generative network models have emerged, including amen (Hoff 2021), bison (Hart et al. 2023), and STRAND (Ross et al. 2023). Here, we contend that these generative network models are the best tools available to move the field of animal social network analysis forward, but for them to be effective, we must integrate non-null models of the sampling processes that generate biased data.

### Empirical data are not easily modeled

Although generative network models provide robust inference by actively estimating and adjusting for correlated random effects at the individual (i.e., node) and dyad level, out-of-the-box implementations of such models should not be expected to solve the problem at hand for researchers in animal behavior. The raw data that researchers have access to are typically affected by sampling and censoring biases that prevent application of most general purpose statistical tools. In an attempt to study the effects of such biases on inferences about animal social behavior, Puga-Gonzalez et al. (2021) applied standard analysis methods to simulated data with biases arising in the data collection process (e.g., the over- or under-sampling of certain categories of individuals, or the censoring of others), and identified false positive rates as high as 61% and false negative rates ranging up to 37%.

Because observational field data collected in natural settings are almost never free from such biases related to sampling and measurement, these findings emphasize a core challenge that needs to be addressed in order to improve the reliability of research findings in the animal networks literature. It is, however, of comfort, that similar concerns about methodological validity have arisen in other sub-fields of network science (e.g., Butts 2003)—and these concerns have been largely resolved by Bayesian approaches.

The gold standard solution to the problem of drawing valid inferences from data that were generated under various forms of measurement bias is simple: we first build a joint generative model of the true biological phenomenon of interest and the bias-inducing measurement process, then we use probability theory (Jaynes 2003)—i.e., Bayesian inversion—to infer the parameters of both the biological phenomenon and the measurement process from observed data (Ross et al. 2023). This approach requires propagating uncertainty and causal effects through multiple simultaneous equations (McElreath 2018), rather than constructing *ad hoc* indices which erase information (e.g., dyad-level sample-size variation) relevant to accurate parameter estimation in subsequent modeling steps.

In what remains of the paper, we will first discuss the prevailing approaches in the field, comment briefly on the known shortcomings of those approaches, and then introduce a formal Bayesian model that rectifies those shortcomings. We analyze simulated data and validate our Bayesian model by demonstrating that the we can recover the parameters of the true social network, even when there are substantial biases in sampling or censoring intensity as a function of individual-level covariates that also affect the structure of the network. We then compare our model to a suite of other approaches using simulated data, demonstrating that the Bayesian approach outperforms all other approaches. All code needed to reproduce the analyses in this paper is available at: https://github.com/SebastianSosa/ASNA_reliability. Additionally, to make application of our methods easy for end-users, we implement our measurement error models in the STRAND R package and provide a full tutorial on data simulation and empirical analysis on the package’s GitHub page: https://github.com/ctross/STRAND.

### Extant approaches for dealing with measurement biases

In animal social network analysis there are by convention two hypothesis testing protocol steps: 1) estimating social interaction patterns among individuals, and 2) testing statistical hypotheses about these patterns. We will outline these steps and provide some commentary on their advantages and disadvantages from a Bayesian perspective.

#### Quantifying interactions

As the first step of an analysis researchers typically calculate a measure of the tendency for individuals to associate (in undirected behavior) or interact (in directed behavior). This measure is referred to as a ‘social index’ and is computed for each pair of individuals (i.e., each dyad). These values are used to create a social network, where each individual is a node and the social index values represent the strength of the connections (edge weights) between nodes. In addition, researchers typically design social indices with the goal of controllingfor sampling effort (i.e., heterogeneity in sampling intensity between individuals).

Currently two main types of social indices are used: association indices (Hubalek 1982; Sailer and Gaulin 1984) and interaction indices. Association indices have been used by behavioral ecologists to estimate the proportion of time that a pair of individuals spends together (Whitehead 2008). The higher the index value, the stronger the dyadic association. The most widely used association index is the simple ratio index (SRI; Eq. 1), which is designed for data collected in discrete sampling periods (e.g., when employing the “gambit of the group”, or scan sampling). The Interaction Index (II; Eq. 2) is primarily utilized by primatologists for data collected during continuous sampling periods, such as during focal-follow sampling. This index estimates the rate of social interactions per unit of time (e.g., the total time of focus). Again, a higher index value indicates a stronger dyadic rate of interaction.

Mathematically, the SRI, Λ_[*i*,*j*]_, is defined as:

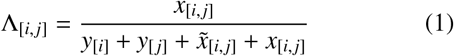

where *x*_[*i*,*j*]_ is the number of sampling periods with observed *i* → *j* ties, *y*_[*i*]_ is the number of sampling periods with only *i* identified, *y*_[*j*]_ is the number of sampling periods with only *j* identified, and 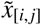 is the number of sampling periods with *i* and *j* identified, but without any *i* → *j* ties.

The Π_[i,j]_ is defined more simply as:

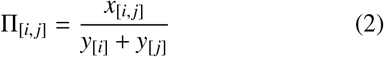

where *x*_[*i*,*j*]_ represents the overall count of *i* → *j* interactions or the total dyadic interaction time, and *y*_[*i*]_ and *y*_[*j*]_ denote the number of observations or the total observation time for individuals *i* and *j*, respectively. The underlying assumptions here are: 1) that *i*→ *j* ties could be detected either during the time that *i* is focal followed, *y*_[*i*]_, or during the time that *j* is focal followed, *y*_[*j*]_, and 2) that the focal follows of individuals *i* and *j* are disjoint (i.e., non-simultaneous), so as to avoid double counting the same ties.

#### A methodological critique

By performing the calculations shown in Eqs. 1 and 2, and dividing by sample size, researchers make all resulting downstream analyses/calculations mathematically fallacious and incapable of supporting any rigorous scientific conclusions. As such, the use of these indices should be avoided, *especially when there is variability in sampling intensity across dyads*. Although many past publications have supported the use of these indices, and many more publications have used them to test scientific questions, the noise and bias introduced by the method is obvious (see Fig. 1). These disadvantages, however, are easily rectified by using Bayesian analogs to the SRI and II.

**Figure 1.**
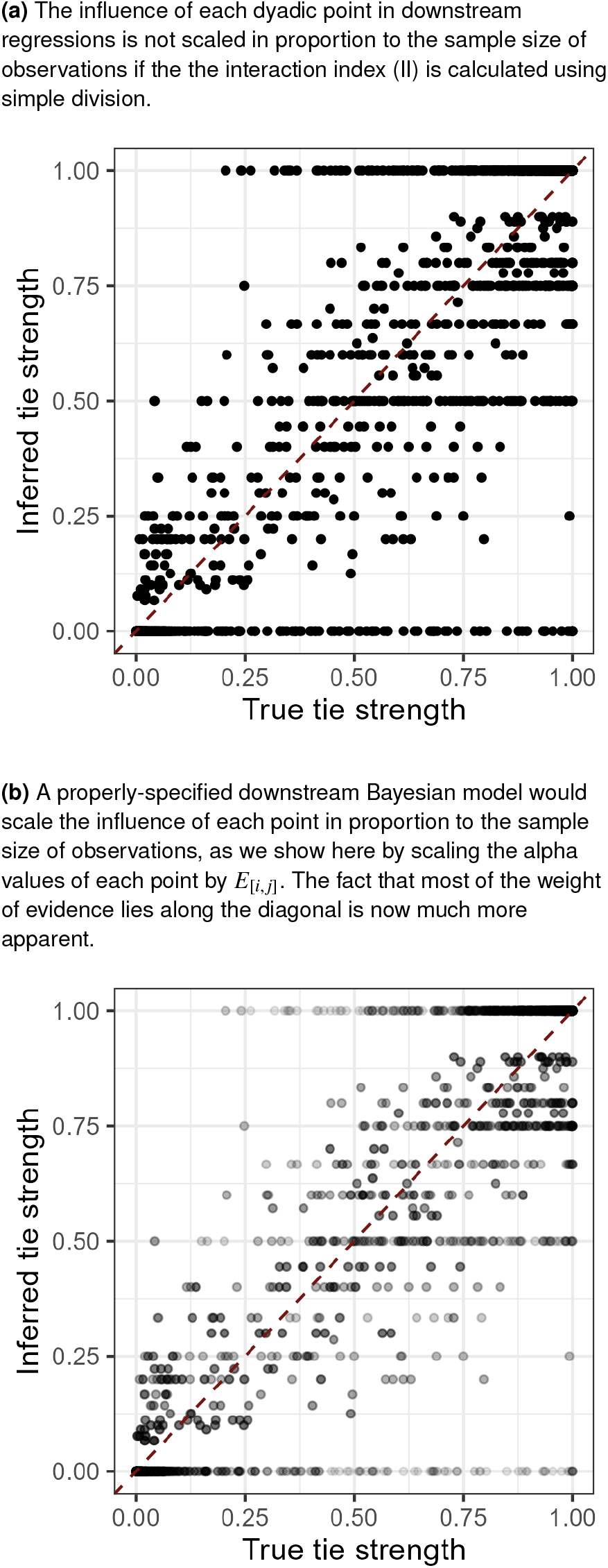
A problem with sociality indices. Example data on tie strength were generated under a model where a covariate *C* has a positive effect on tie probability, ϕ_[i,j]_, but a negative effect on sampling effort, *E*_[i,j]_. We plot the true dyadic tie strength on the x-axis and the interaction index, II, on the y-axis. In frame (a), we see that many points lie along the diagonal (as expected), but there are also strips of points in horizontal lines at rational numbers with small denominators: e.g., at 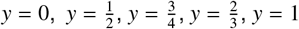. Due to the small sample sizes for points on these bands, the inferred tie strength is not that reflective of true tie strength. Simply plugging in Interaction Index estimates into downstream regressions thus leads to poor statistical inference, because the dyadic samples based on few observations obscure the signal in the rest of the data.

This being said, the *motivation* for the SRI and II is clearly well-founded. The raw count or duration of interactions, *x*_[*i*,*j*]_, is analytically meaningless without corresponding information for each dyad that quantifies the *opportunity* or *sampling risk, E*_[*i*,*j*]_, for dyadic ties to be observed in the first place. This is especially true when there is heterogeneity in sampling intensity across individuals or dyads.

Well-specified Bayesian models using the II approach, for example, could be written by letting *E*_[*i*,*j*]_ = *y*_[*i*]_ + *y*_[*j*]_, and then using probabilistic models like:

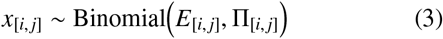

to infer the full posterior distribution for each Π_[i,j]_, rather than using the division operator to get a simple point estimate, which discards information on the precision/certainty of that point estimate. If sample size varies across dyads, then construction of an index using Eqs. 1 or 2 means that all downstream statistical analysis depending on that index will violate the laws of probability theory^*^, and will therefore be unreliable. In a Bayesian analysis, however, Eq. 3 can be fit jointly with a downstream model that uses the Π_[i,j]_ parameter array as either a predictor variable, or an outcome variable. That downstream model would be well defined and consistent with the laws of probability theory, because all posterior uncertainty in each element of Π_[i,j]_ is propagated and thus accounted for in the downstream analysis. If only the point estimates of Π_[i,j]_ are produced using Eqs. 1 or 2, then serious methodological problems can—and do routinely—arise, and these problems will be strongest precisely when *when there is variability and*/*or bias in sampling intensity across dyads*, which is unfortunately the exact same context in which *control for variability*/*bias in sampling is most important*. Dividing by sample size causes the weakest data points—with the highest sampling-error-to-signal-ratio—to have disproportionate leverage on downstream parameter estimates. For a visualization of this issue see Fig. 1.

#### Testing for patterns

As the second analytical step, researcher typically test statistical hypotheses about social interaction patterns among individuals by computing various node-based measures from point-estimates of the network of associations. For example, they might sum rows of an adjacency matrix to get an estimate of out-degree (for binary networks) or out-strength (for weighted networks). These metrics measure the tendency of individual *i* to act on others. However, using such aggregated measures to test hypotheses about individuals is not ideal because features of each individual *j* influence the row sums for each individual *i*, and likewise when calculating column sums to estimate in-degree or in-strength (i.e., the tendency of individual *i* to be acted upon by others). This interdependence violates the assumptions of common parametric tests. Moreover, each association/interaction is counted twice, once for each individual in the dyad, which further violates the assumptions of common parametric tests. Therefore, much methodological work in the animal social network literature has focused on developing techniques to enable valid hypothesis testing in spite of these interdependencies.

Permutation methods have been the standard tools and they come in two forms: network permutations and pre-network permutations (Farine 2013). For more detailed discussions of the principles of permutation approaches readers can see: Farine (2013) and Sosa et al. (2021b). Issues related to unacceptable false positive and false negative rates were originally demonstrated in workflows that used either network permutations or pre-network permutations (Puga-Gonzalez et al. 2021; Hart et al. 2022). To address these issues, double permutation workflows, which combine both approaches, have been developed and tested on biased data. Double permutation approaches appear to provide better control of false positives and false negatives than other permutation approaches (Farine and Carter 2022). However, they represent an alternative analytical pipeline to the Bayesian approach we advocate here and so we do not discuss them further.

#### A methodological critique

Permutation methods have long faced criticism on purely logical grounds (see: Krackhardt 1992). Moreover, researchers generally care about *the probability of some biological phenomenon conditional on the actually observed data* (which requires a Bayesian analysis), rather than the probability of data that was never actually observed under a causally irrelevant null model (*which is what a permutation test provides*). In our opinion, however, the biggest problem with permutation methods in applied settings is that end-users want to use them to account for the interdependencies in network data, but they simply do not do this (Hart et al. 2022). Debates on their use, however, distract from the gold-standard approach of using correlated random effects to rigorously account for such interdependencies. Here, we follow the guidance of network scientists and applied statisticians (Snijders and Kenny 1999) and advocate for the use of highly parameterized, generative network models, like the Social Relations Model (Kenny and La Voie 1984) and its many extensions (Back and Kenny 2010; Christensen et al. 2003; Eichelsheim et al. 2009; Koster and Leckie 2014; DeTroy et al. 2021; Redhead et al. 2023a; Pisor et al. 2020). Across several disciplines, these models have been used to control for network interdependencies, while simultaneously permitting parametric estimation of key network properties, like block structure, nodal heterogeneity, generalized reciprocity, dyadic reciprocity, and higher order multiplex reciprocity structures (Redhead et al. 2024).

### Advancing the state of network science in animal behavior

The studies that first highlighted problems with permutation approaches and proposed new methods (e.g., Weiss et al. 2021; Puga-Gonzalez et al. 2021; Beaulieu et al.) have been employing the same simulation protocol—demonstrated in Appendix 1. This simulation has some issues of its own. First, these simulations were designed to generate a “bias of interaction”. This bias pertains to failing to detect key associations or interactions, *while observing* key individuals. It differs from “sampling bias”, which refers to heterogeneity in the amount of time individuals are observed. Interaction bias is thus a form of data censoring, where the researcher does not know that he or she is failing to note true interactions that did occur. In contrast, observation bias is known to the observer and is thus easier to account for. However, choices in the coding of the simulation protocol inadvertently lead to the simultaneous creation of both observation bias and interaction bias. This complicates separation of the effects of each form of bias.

To tackle these concerns we developed a new simulation protocol—included in the `STRAND` package—which allows users to independently specify sampling biases and interaction/censoring biases. Using this simulation, we created various scenarios to assess the reliability of results obtained through different methods including interaction indices, network permutations, double permutations, Bayesian generative networks, and a new Bayesian measurement error model that will be introduced in the methodology section.

In this paper, we have multiple goals. First, we aim to assess whether issues related to false positives and false negatives in animal social network analysis methods are associated more with observation or interaction biases. We are expecting to observe accurate results for all Bayesian approaches regarding “bias of observation”, but are expecting methods that divide by sample size to perform comparatively poorly. We also expect that once “interaction bias” is introduced in the simulation, all methods—other than our new Bayesian measurement error model—will be inaccurate, as none of the other protocols structurally account for this type of bias. Our secondary objective is to determine how analyses based on Bayesian generative networks compare with extant methods in animal social network analysis. Over the years, multiple approaches have been proposed. Researchers faced with such a plethora of methods may encounter difficulties in assessing the strengths and weaknesses of each. The present study will provide clear performance benchmarks.

## Methods

### Modeling animal networks with sampling biases and censoring issues

In the vast majority of cases, researchers in animal behavior collect network data in the form of numerical outcomes— i.e., the number of times that interactions *were observed*, conditional on the number of times that interactions *could have been observed*. For example, researchers may conduct a specific number of observations (scans or focal follows) and the outcome data, *Y*_[i,j]_, might reflect the number of observations in which directed (e.g., grooming or aggression events) or undirected (e.g., spatial associations) interactions between individuals *i* and *j* were observed. The number of observations in which interactions could have been detected for a given dyad is called the sampling effort, *E*_[i,j]_. Sampling effort is an *exposure* variable, which strongly—indeed, *proportionally*—influences the outcome counts. To account for variation in sampling effort across dyads in a way that fully propagates uncertainty (see Ross et al. 2023, for derivation), a Bayesian model of network data will generally take the form:

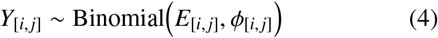

where *ϕ*_[i,j]_ is the latent directed connection strength between *i* and *j*, represented as the probability that a directed tie from *i* to *j* occurs in a given observation period. *Y*_[i,j]_ is typically a count of observation periods in which a tie was observed, and *E*_[i,j]_ is the number of observation periods in which ties between *i* and *j* could have been detected. Note that, in the generative context, *ϕ* represents a true biological phenomenon, while the outcome variable, *Y*, and the sampling variable, *E*, may be strongly affected by the sampling protocol, researcher behavior, and other features of animals *i* and *j* that might interfere with observation (e.g., cryptic coloration).

The goal of analysis is to measure *Y* and *E* and then recover *ϕ*. This can be complicated, however, if there are structural factors that influence both *ϕ* and *Y* and/or *E*. Here we will consider two potential causes of bias: (1) sampling bias, where features of *i* and *j* influence *E*_[i,j]_, and (2) censoring, where features of *i* and *j* influence measurement of *Y*_[i,j]_ given *E*_[i,j]_, independent of *ϕ*_[i,j]_.

We present a full Bayesian model of the data generating process that includes both the “true”, biologically meaningful, weighted network of dyadic interactions, and the process of measurement that might produce biased outcome measures. We start by describing the sub-model underlying the true network and then integrate the sub-models underlying the measurement process. We conclude by showing how proper model specification permits the recovery of the true social network from biased measurements.

### The Social Relations Model (SRM)

#### Model definition

We begin the process of constructing the model described in Eq. 4 by providing a generative model for the *true* latent directed connection strength between individuals *i* and *j, ϕ*_[i,j]_. In order to generate networks with empirically plausible typologies, it is generally necessary to define a model for *ϕ*_[i,j]_ that includes correlated random effects for the propensity to initiate and receive interactions (i.e., nodal random effects), and correlated dyad-level random effects for the propensity of *i* to send to *j* and *j* to send to *i*. Additionally, such models should permit inclusion of covariate effects that influence block/group structure, node-level tie propensity, and dyad-level tie propensity. See Fig. 2 for an example network generated under such a model.

**Figure 2.**
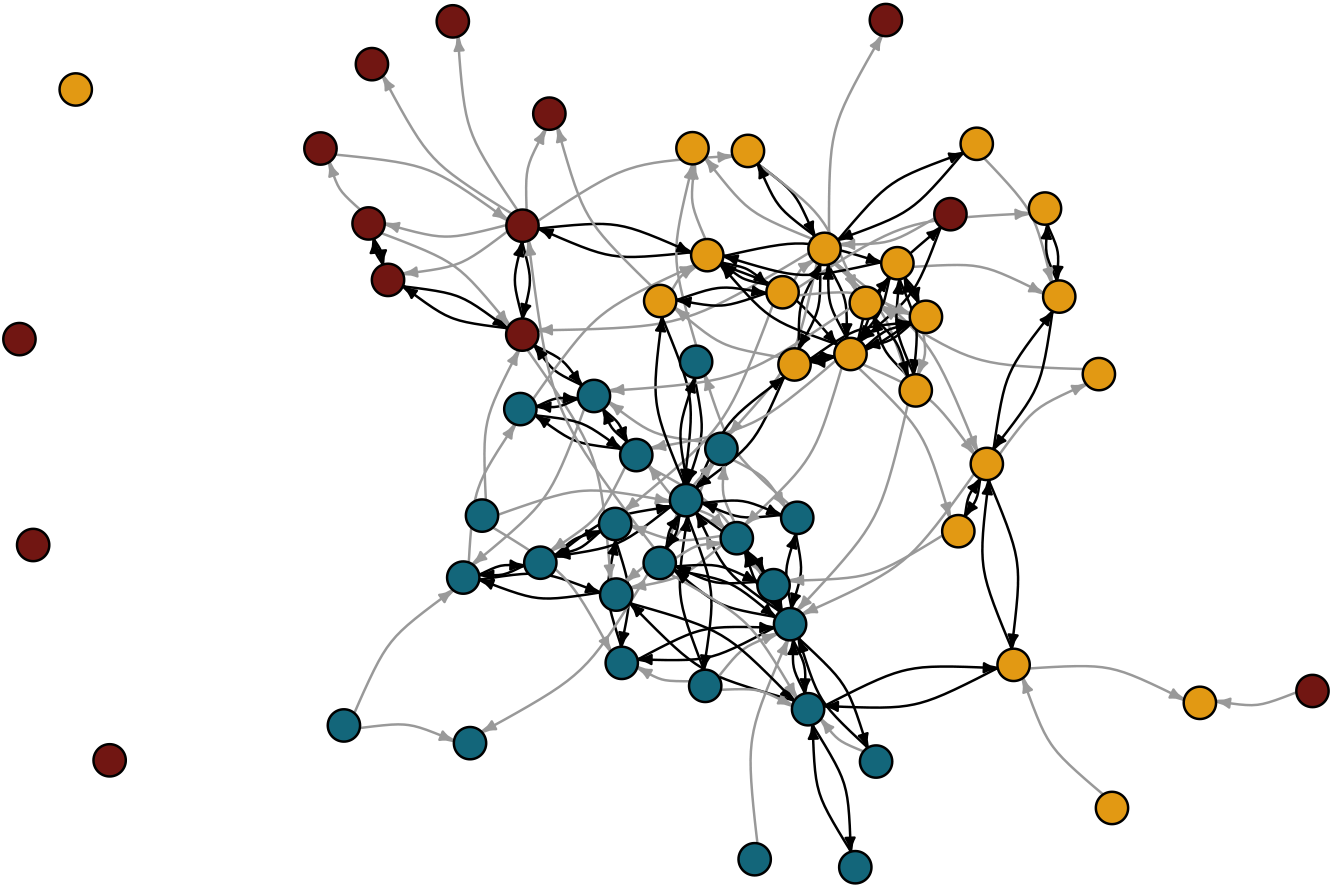
Simulated animal network data. Data were simulated using the generative model described in Ross et al. (2023). Nodes are colored by group. The stochastic block model introduces gross substructure, where ties within groups are more likely than ties between groups. These rates can be varied continuously, producing networks with no group structure on one extreme, to networks fully segmented by group identity on the other. Random effects for sending and receiving ties can lead to heterogeneity in number of social partners (i.e. degree). Here, some individuals have small degree and are connected to the network by only a single tie, and others have a high degree and are connected to many other individuals. As with the parameters controlling group structure, the parameters controlling variance and covariance in individual-level propensity to send and receive ties can be varied continuously, producing networks with approximately uniform degree on one extreme, to networks with highly unequal degree distributions on the other extreme. Finally, we note that some dyads form reciprocal ties (note the black, bidirectional arrows), while other dyads are linked only by unidirectional ties (note the grey, unidirectional arrows). The parameters controlling variance and covariance in dyadic random effects are continuously variable and can produce networks with close to no dyadic tie reciprocation on one extreme, and networks with high rates of tie reciprocation on the other. Through Bayesian inversion, we can infer the parameters of the model used to generate this network from the dyad-level interaction data; this allows us to learn how individual and relational attributes are related to various aspects of social network structure.

Following prior work in Ross et al. (2023) and Redhead et al. (2023b), we recommend the use of the social relations model (Kenny and La Voie 1984; Snijders and Kenny 1999; Back and Kenny 2010) with an additional set of stochastic blockmodel parameters (Holland et al. 1983; Karrer and Newman 2011; Peixoto 2019) to account for group structure when necessary. As such, *ϕ*_[i,j]_ can be modeled as:

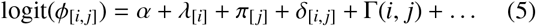

where *α* is an intercept term, *λ* is a vector of individual-specific focal/sender/outflow effects, *π* is a vector of individual specific target/receiver/inflow effects, *δ* is a matrix of dyadic effects, Γ(*i, j*) is a function giving a dyadic intercept offset as a function of group/block structuring variables, and the ellipsis signifies any linear model of coefficients and focal, target, or dyadic covariates.

For example, if *C* is an animal-specific measure, like coloration, and *Q* is a dyad-specific measure, like a matrix of genetic relatedness, then the ellipsis may be replaced with: *κ*_[1]_*C*_[*i*]_ + *κ*_[2]_*C*_[ *j*]_ + *κ*_[3]_*Q*_[i,j]_, to give the effects of coloration on the probability of sending ties to any target and receiving ties from any target, and the effect of kinship on the probability of dyadic ties.

To model block structure, we can consider a list of *V* categorical variables—e.g., sex or group—describing individuals *i* and *j*. Let *B*_[*v*,*u*,*w*]_ be a three-dimensional parameter array where *v* runs over variables and *u* and *w* run over the category/block levels within variables. Finally, let the function *b*(*i, v*) return the block of individual *i* for variable *v*. Then we can define Γ(*i, j*) such that:

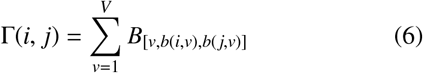

where the probability of a tie from individual *i* in block *b*(*i, v*) to individual *j* in block *b*( *j, v*) for variable *v* is controlled by the corresponding entry in the array of block parameters, *B*_[*v*,*b*(*i*,*v*),*b*( *j*,*v*)]_. From this set of block parameters we can calculate any of a variety of assortativity coefficients (e.g., Newman 2003), which describe how much individuals tend to associate with others of the same type as themselves.

#### Priors

To complete the model definition, we use weak priors. We model the sender and receiver effects jointly using a multivariate normal distribution. This allows for generalized correlations at the individual level to be detected— i.e., we can detect if individuals who generally groom others are also more likely to be groomed in general by others. For computational efficiency (Stan Development Team 2021; Lewandowski et al. 2009) it is best to write the multivariate normal as:

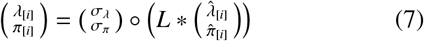

where *L* is a Cholesky factor from the decomposition of the 2 ×2 correlation matrix with *ρ* on the off-diagonal, and 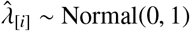 and 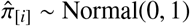 are unit-normal random effects. Weakly informative priors may then be independently specified on the variance and correlation terms (Lewandowski et al. 2009):

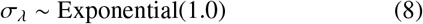

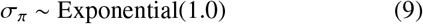

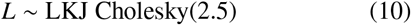

We model the dyad-level random effects jointly using a multivariate normal distribution as well. This allows for dyadic correlations to be detected—i.e., we can detect if the probability of focal *i* grooming individual *j*, increases with the probability that focal *j* grooms alter *i*.

Similar to the model for individual effects, we can write the model for dyadic effects as:

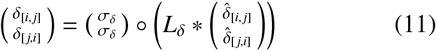

where 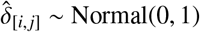 have unit-normal priors and the variance and correlation terms have weakly informative priors:

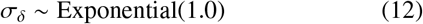

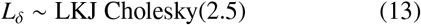

In order to prevent over-fitting in small samples, we recommend standardizing predictor variables and using weakly regularizing priors on the *κ* terms:

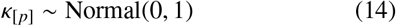

However, these priors and all other default priors discussed below can be modified by STRAND users according to their needs by passing in a labeled list of priors when calling the model function.

Lastly, the diagonal elements of *B*_[*v*]_, which control the frequency of ties within blocks, will generally have slightly higher prior weight than the off-diagonal elements, though other topologies are possible (see Batagelj 1997). For example, we might write:

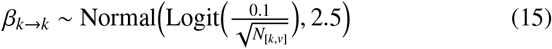

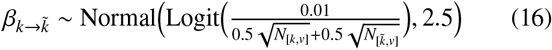

Here, *k* → *k* indicates a diagonal element and 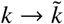 indicates an off-diagonal element. The scalar of 0.1 in Eq. 15 places higher prior density on the diagonal of *B* (which controls the probability of within-block ties), than the off-diagonal of *B* (where the scalar of 0.01 from Eq. 16 generates reduced prior between-block/group tie probability). The scalars of 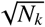 ensure that prior strength scales with sample size at roughly the same rate that we see in empirical data sets (Ready and Power 2021). The standard deviation of 2.5 in both equations causes the overall prior to be quite weak, and thus allows the data to dominate the posterior.

### Sampling bias

The next step in building the model described in Eq. 4 is to provide a generative model for the sampling effort between individuals *i* and *j, E*_[i,j]_. Most animal behavior data collection protocols can be separated in two categories, those based on discrete-time sampling rules and those based on continuous-recording sampling rules (Bateson and Martin 2021). Discrete-time sampling rules are instantaneous or zero-one sampling designs, such as the “gambit of the group” or “scan sampling” protocols (see: Sosa et al. 2021b, for further details). These approaches involve recording the count of behavioral events but not their duration. In such data collection protocols, the sampling effort directed toward individual *i* would be the count of sampling periods during which it was possible to observe ties involving individual *i*. Continuous-recording sampling designs instead measure the temporal duration of behavioral states—e.g., how many seconds of grooming occurred during a follow, or the length of time that two GPS trackers were within a threshold distance of each other (He et al. 2023). These designs also measure rates of interactions per unit of time— e.g., how many aggressive interactions occur between two individuals during a follow. The sampling effort directed toward individual *i* would be the summed recording time of all behaviors (including no-behavior/rest) produced by individual *i*. Whether we focus on discrete or continuous sampling designs^†^, a simple model of sampling effort can be written as:

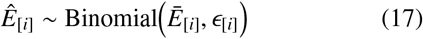

Here, *Ē* _[*i*]_ refers to maximum possible sampling effort (i.e., the total number of scans a researcher dedicated to following individual *i*, or the total amount of time a GPS tracker was placed on individual *i*). Likewise, *Ê*_[*i*]_ is the realized sampling effort for individual *i* which may differ, for example, if some individuals evade the researcher or take refuge in areas where the GPS trackers cannot record accurately. Lastly, *ϵ*_[*i*]_ is the probability that individual *i* is actually observed by the researcher during a given sampling time-point. We allow the parameter *ϵ*_[*i*]_ to vary as a function of individual-level characteristics that may also influence network structure. For example, an animal with a cryptic phenotype may be more likely to be unobserved—even as the focal in a focal-follow—if their coloration allows them to escape the attention of the researcher performing behavioral observation. Similarly, individuals with a cryptic phenotype may be under-sampled, because they are less likely to be found on a given day and are thus subject to fewer focal follows than desired overall.

We formalize this by modeling:

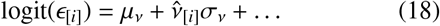

where *μ*_*ν*_ is an intercept, *σ*_*ν*_ is a scalar for the variance of random effects, 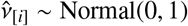, and the ellipsis signifies any linear model of coefficients and individual-level covariates. For example, if *C* is an animal-specific measure, like a binary variable for cryptic coloration, then the ellipsis may be replaced with: *κ*_[4]_*C*_[*i*]_, to give the effects of coloration on the number of sampling events.

Using the interaction index method discussed above, the dyad-level sampling effort is given by the formula:

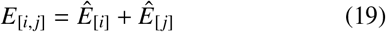

as ties between *i* and *j* can be detected either when *i* is subject to observation or when *j* is subject to observation. It should be noted that Eq. 19 is only a good approximation if it is true that *i → j* ties could be recorded if either *i* or *j* is the focal individual being observed.

From our formalization in Eq. 4, it is clear that variation in sampling effort should have little to no effect on our ability to accurately identify *ϕ*_[i,j]_ and downstream quantities, since a binomial model of *Y*_[i,j]_ given *E*_[i,j]_ provides information about the posterior value of *ϕ*_[i,j]_ that does not further depend on *E*_[i,j]_; however, the narrowness of the posterior distribution of *ϕ*_[i,j]_ does depend on *E*_[i,j]_, as we can be more confident in estimates of *ϕ*_[i,j]_ that are based on more observations.

### Interaction bias, or censoring

Censoring, or interaction bias, is a more severe problem. In this case a researcher might perform *E*_[i,j]_ observations and detect *Ŷ*_[*i, j*]_ ties. However, if the researcher is observing animal *i* and animal *j* has a cryptic phenotype, then it is possible that ties from *i*→ *j*—or *j* → *i*—did occur, but the researcher did not detect them. For a single observation let the indicator of a true tie be *Q*_[i,j]_ ∈ {0, 1}, the indicator of a detected tie be 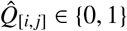, and the indicator for individual *i* being detectable be *D*_[*i*]_ ∈{0, 1} . Under these assumptions we find that:

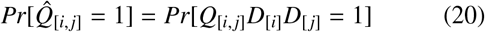

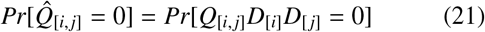

where 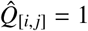 can only occur when both *i* and *j* are detectable and a true tie occurs. 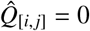, however, occurs as a mixture of three non-disjoint causes: (1) cases where no tie occurs, (2) cases where *i* was not detectable, and (3) cases where *j* was not detectable.

Let *η*_[*i*]_ = *Pr*[*D*_[*i*]_ = 1] describe the detectability of individual *i*, and *ϕ*_[i,j]_ = *Pr*[*Q*_[i,j]_ = 1] describe the true tie probability. We note that Eqs. 20 and 21 define the probability mass function of a Bernouli random variable and so we can aggregate over scans, *E*_[i,j]_, to yield a Binomial model. Thus we can rewrite Eq. 4 as:

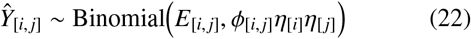

As with the model for sampling time, we can let *η*_[*i*]_ depend on individual-specific covariates. To model the probability of censoring, we can model 1 − *η*_[*i*]_:

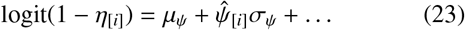

where *μ*_*ψ*_ is an intercept, *σ*_*ψ*_ is a scalar for the variance of random effects, 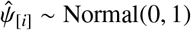, and the ellipsis signifies any linear model of coefficients and individual-level covariates. For example, if *C* is an animal-specific measure, like a binary variable for cryptic coloration, then the ellipsis may be replaced with: *κ*_[5]_*C*_[*i*]_, to give the effects of coloration on censoring probability.

Here, finally, we run up against the limits of inference. Because the terms *ϕ*_[i,j]_ and *η*_[*i*]_ multiply, the likelihood for the network model cannot distinguish the effect of a variable—*C*_[*i*]_, for example—that affects both the odds of a true tie and also affects censoring probability.

The only way to permit accurate inference is to use an independent source of ground-truth data to anchor estimation of *η*_[*i*]_. If, for example, a researcher conducts 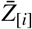 trials to encounter individual *i* in a setting where individual *i* is known to be present and observes individual *i* in *Z*_[*i*]_ of those trials, this permits integration of a sub-model like the following:

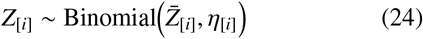

which, in turn, permits estimation of social network parameters that are robust to censoring and interaction bias.

Another possibility to infer *η*_[*i*]_ and *η*_[ *j*]_ involves investigation of the discordance between *i* → *j* and *j*→ *i* tie counts among focal follows in which *i* and *j* are subjects; however, the math required to formalize this model goes beyond the scope of this paper.

### Validating the STRAND Binomial SRM with measurement error

In order to validate the model outlined above, we code it first as a data simulation function in R using STRAND and second as a Bayesian inference model. We then simulate network data—with either sampling bias or censoring bias—as a function of a covariate that also affects the propensity of individuals to act towards others (i.e., an out-strength effect) or be acted upon by others (i.e., an in-strength effect). We then ensure that our inferential model recovers the values of the parameters used to simulate the data. In Fig. 3, we simulate *n* = 22 network datasets subject to either bias in sampling (*n* = 11) or censoring (*n* = 11) and compare two analysis models on each. The first, plotted in blue, is the standard binomial network model included in STRAND (Ross et al. 2023). This model assumes— counterfactually in this case—that the collected data represent “true” information, without any measurement bias. The second model, plotted in goldenrod, is the binomial network model introduced above, which formally accounts for sampling and censoring biases. For each model, each of the 11 datasets was created assuming that an individual-level predictor variable, *C*, had a strong positive effect on individual emitted interactions (i.e., out-strength) of 1.9, and a moderate positive effect on individual received interactions (i.e., in-strength) of 0.55. These ground-truth values are depicted as dashed horizontal lines. In each dataset, that same variable, *C*, had an effect on the log-odds of the sampling rate or censoring level, across datasets we let these effects range from -4.5 to 4.5. In the top panels of Fig. 3, we see only goldenrod curves, because only the new model has parameters to estimate the predictors of measurement bias. Across datasets, the new measurement-error models correctly estimate the level of sampling or censoring bias, which we can see by the dashed diagonal line falling within the 95% credible region.

**Figure 3.**
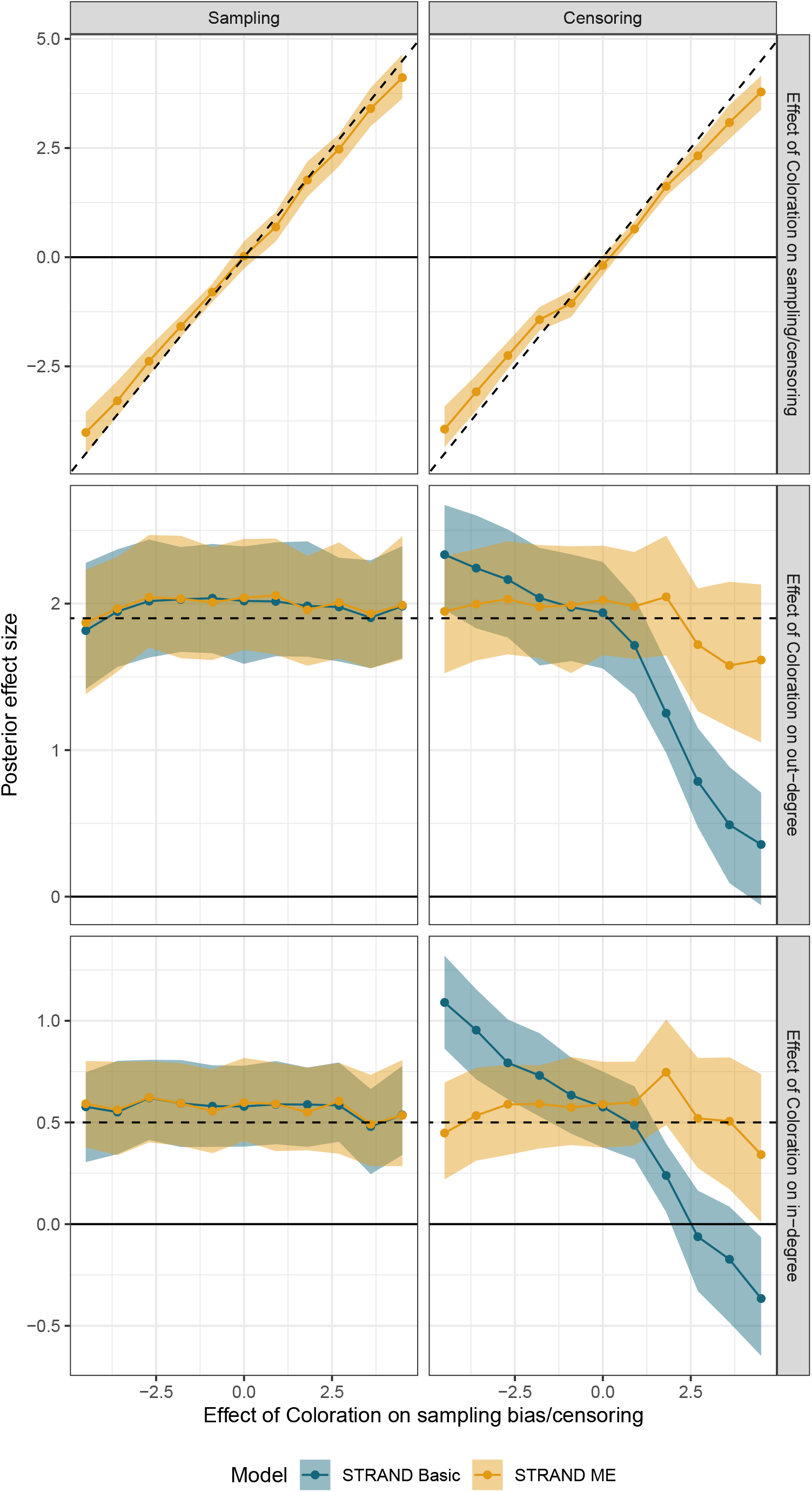
Parameter recovery via the analysis of simulated datasets. The basic binomial network model from STRAND is plotted in blue and the new binomial model which simultaneously accounts for measurement biases is plotted in goldenrod. For interpretation of results, see discussion in the main text. The takeaway message is that the standard binomial model is sufficient to deal with biases in sampling effort, but only the improved model can accurately recover parameters if the data are subject to censoring (i.e., interaction bias).

With respect to sampling bias, we find that the basic binomial network model in `STRAND` is fully robust. That is, in the left-hand panels of Fig. 3, the basic binomial model and the binomial model which accounts for measurement biases, yield identical parameter estimates. This is expected behavior because the binomial model naturally controls for sampling heterogeneity appropriately by itself. However, with respect to censoring bias plotted in the right-hand panels, we find that the basic binomial network model in `STRAND` is not robust. If censoring bias is extreme in either direction, then the basic binomial model (plotted in blue) is unable to accurately recover the effects of *C* on individual in-degree and out-degree. However, applying the new model (plotted in goldenrod) to the same data, we are able to accurately recover the true effects of *C* on individual out-strength and in-strength.

### Testing animal social network analysis methods

#### Scenarios

Using the simulation model explained earlier, we create three scenarios. In scenario 1, we assess the ability of various animal social network analysis methods to accurately infer the effects of two variables, *coloration* and *shyness*, on nodal out-strength and in-strength when no biases in sampling intensity or censoring probability are present. Across simulations, we vary only the effect of coloration on nodal out-strength (from -2.0 to 2.0) while keeping all other parameters fixed. In particular, the true effect of coloration on nodal in-strength is set to a moderate value of 0.55. The shyness variable is set to be weakly correlated with coloration, but the effect of shyness on nodal out-strength and in-strength is set to be zero. Thus, our simulation allows us to test for false positives and false negatives, over a range of values for the effect of coloration on nodal out-strength.

In scenario 2, we examine the reliability of various methods when sampling bias is introduced. In this scenario, we keep all parameter settings from scenario 1, but we set the effect of coloration on sampling intensity to be -2.75. This leads to a strong masking effect, where individuals with high levels of coloration are less likely to be observed, but the researcher is aware of the variation in dyadic sample size induced by this effect.

In scenario 3, we examine the reliability of various methods when censoring is introduced. In this scenario, we keep all parameter settings from scenario 1, but we set the effect of coloration on censoring probability to be 2.75. This leads to a strong masking effect, where ties emitted by individuals with high levels of coloration are less likely to be observed, but the researcher perceives little variation in dyadic sample size.

#### Tested approaches

For each simulation, we evaluated eight different analysis routines: (1) the standard `lm` function in base R, (2) the `ame` function from the `amen` package developed by Hoff (2021), (3) the `mrqap.dsp` function from the `asnipe` package developed by Farine and Whitehead (2015b), (4) the `bison_model` function from the `bisonR` function developed by Hart et al. (2023), (5) the `fit_block_plus_social_relations_model` function from the `STRAND` package developed by Ross et al. (2023), and (6) the `fit_block_plus_social_relations_model me` function from the `STRAND` package, developed specially for this paper. Additionally, we used the `ANTs` package developed by Sosa et al. (2020) to implement two further analyses. In method (7), we use `ANTs` to conduct a simple nodal regression based on the interaction index. For method (8), we use `ANTs` to conduct the same analysis as before, but using permutation methods to calculate *p*-values.

For each approach, in each simulation, we calculate the estimated effect size for each variable of interest, along with the relevant *p*-values^‡^. Although we advocate for the use of methods which permit quantitative recovery of effect size estimates, many of the methods that we compare, do not— even in principle—permit such parameter recovery, forcing us to compare *p*-values.

## Results

### Scenario 1

In scenario 1, we simulate network data with no biases in sampling effort and no censoring issues. We simulate 11 datasets, holding all parameters constant across simulations except the effect of coloration on out-strength, which we denote as *κ*. We explore values of *κ* ∈ ( −2.0, 2.0). We compare analysis methods with a focus on assessing if each method: (1) correctly inferred the effect of coloration on out-strength (true values are given by *κ*), (2) correctly inferred the effect of coloration on in-strength (true value is 0.55, a weak effect), (3) correctly inferred that there is no true effect of shyness on out-strength, and (4) correctly inferred that there is no true effect of shyness on in-strength.

The first panel of Fig. 4 shows that all models correctly detect the effect of coloration on out-strength. As expected, all methods have peak *p*-values at values of *κ* = 0, which rapidly drop below the threshold of 0.05 as *κ* increases towards 2.0 or decreases towards -2.0. The second panel of 4 shows that all methods accurately infer a significant effect of coloration on in-strength for values of *κ* < 0. However, when *κ* > 0, permutation based approaches (i.e., MRQAP from `asnipe`, node label permutations from `ANTs` ) and linear regressions without permutations fail to detect true effects of coloration on in-strength, whereas the Bayesian models continue to recover the true effect of coloration on in-strength over the whole range of *κ*. The third panel of Fig. 4 shows that most methods perform well across the whole parameter range: shyness has no effect on out-strength, and most methods return large *p*-values. For high values of *κ*, however, a few methods (`ANTs Rates Permuted, lm Rates`, and `AMEN`) yield small, spurious *p*-values. The fourth panel of Fig. 4 shows that all models correctly indicate that there is no effect of shyness on in-strength.

**Figure 4.**
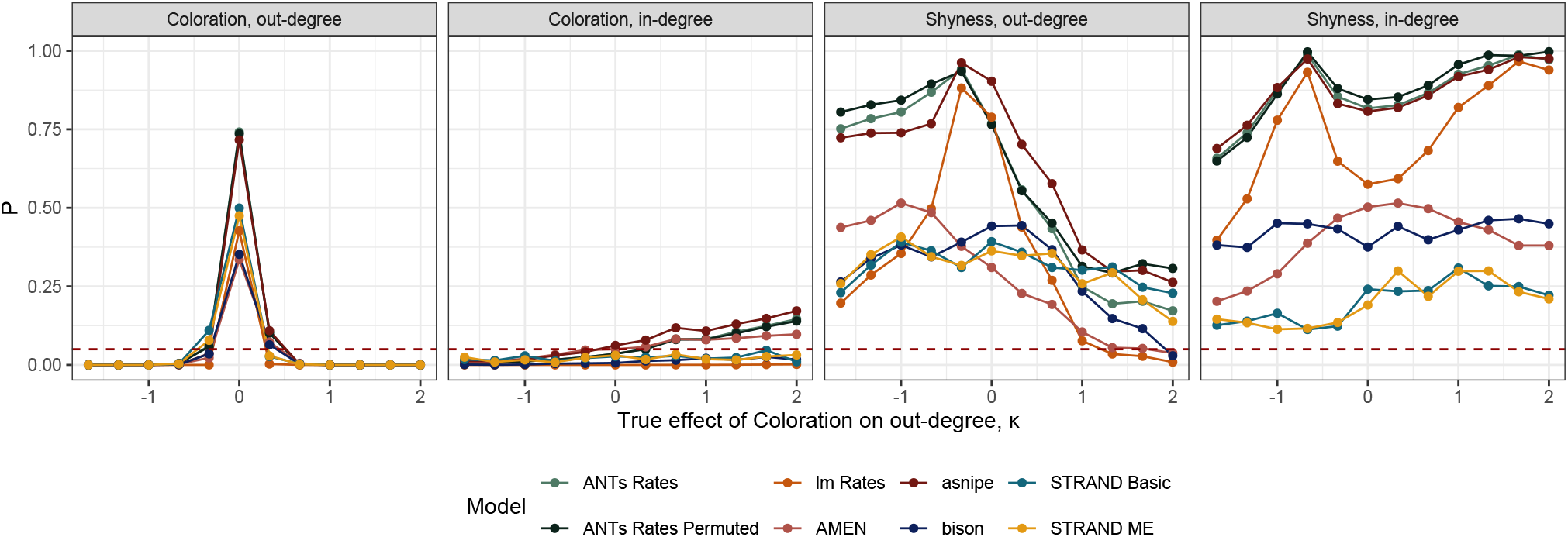
Analysis of simulated datasets with no sampling bias or censoring, using eight different network approaches. In each panel, we plot *p*-values for the indicated coefficient estimates on the y-axis, and the true effect of coloration on out-strength (κ) used in the simulation on the x-axis. If a model is performing correctly, then we should observe the following four things: (1) In the first panel, the *p*-values should be close to zero when κ is either very large or very small, and the *p*-values should be large when κ is close to zero. In other words, conditional on our sample size of *n* = 65 individuals, we should be able to detect large true effects, but not infer significant effects when none exist. (2) In the second panel, *p*-values should always be small, because coloration was set to have a positive effect on in-strength in all simulations. (3) In the third panel, *p*-values should always be large, because shyness has no effect on out-strength in the generative model. In other words, if a model is performing well, it should not infer a significant effect of shyness when such an effect does not exist. (4) In the fourth panel, *p*-values should always be large because shyness has no effect on in-strength. In terms of performance, all models correctly detect the effect of coloration on out-strength in panel one. All methods have peak *p*-values at values of κ = 0, which rapidly drop below the threshold of 0.05 as κ increases towards 2.0 or decreases towards -2.0. In panel two, all methods accurately recover the significant effect of coloration on in-strength for values of κ < 0. However, only the Bayesian models recover the significant effect of coloration on in-strength for values of κ > 0. In panel three, all methods perform well across the whole parameter range. Shyness has no effect on out-strength, and all methods return large *p*-values. In panel four, all models correctly indicate that there in no effect of shyness on in-strength.

### Scenario 2

In scenario 2, we simulate network data with biases in sampling effort that are caused by a variable, coloration, that also affects out-strength and in-strength. As in scenario 1, we simulate 11 datasets, holding all parameters constant across simulations except the effect of coloration on out-strength.

The first panel of Fig. 5 shows that most models correctly detect the effect of coloration on out-strength. Most methods have peak *p*-values when *κ* = 0, which rapidly drop below the threshold of 0.05 as *κ* increases towards 2.0 or decreases towards -2.0. The second panel of Fig. 5, however, shows that several methods infer significant effect of coloration on in-strength only for values of *κ* < − 1. Here, it becomes obvious that the permutation methods common to animal network analysis do not permit accurate inference in the presence of biases in sampling intensity. In contrast, only `BISON` and `STRAND`, recover the true effect of coloration on in-strength for all values of *κ*. The third panel of Fig. 5 shows that all methods perform well across most of the parameter range: shyness has no effect on out-strength, and most methods return large *p*-values however, a few methods (`lm Rates, AMEN, BISON`) yield small, spurious *p*-values for values of *κ* > 1. The fourth panel of Fig. 5 shows that all models correctly indicate that there in no effect of shyness on in-strength.

**Figure 5.**
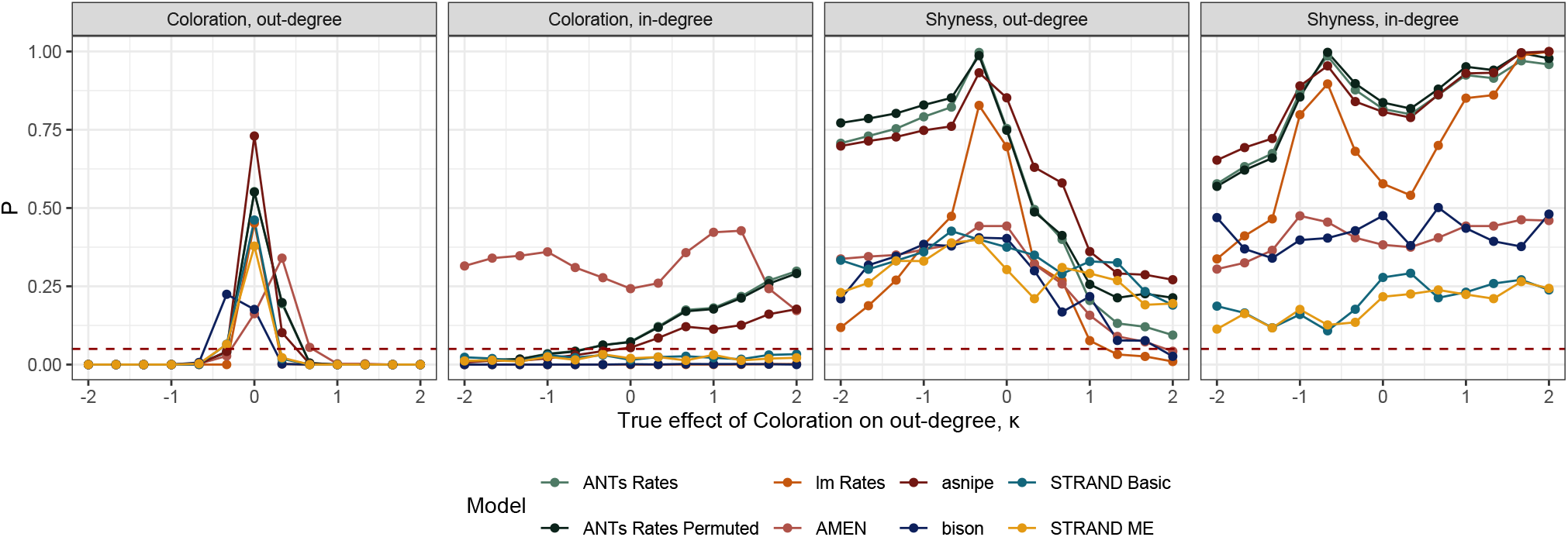
Analysis of simulated datasets with sampling bias, using eight different network approaches. See the caption to Fig. 4, for a description of how to interpret each panel below. In terms of performance, most methods permit accurate estimation of the effect of coloration on out-strength in panel one. Most methods have peak *p*-values at values of κ = 0, and rapidly drop below the threshold of 0.05 as κ increases towards 2.0 or decreases towards -2.0. In panel two, only the Bayesian models accurately detect the true effect of coloration on in-strength. The other methods fail for κ > −1, likely because the effect of coloration is only moderate in strength and construction of the sociality index divides out sample size information, obscuring the true effect. In panels three and four, all methods perform well across the whole parameter range. All models correctly indicate that there is no effect of shyness on out-strength or in-strength.

### Scenario 3

In scenario 3, we simulate network data with censoring effects caused by a variable, coloration, that also affects out-strength and in-strength. As in scenarios 1 and 2, we simulate 11 datasets, holding all parameters constant across simulations except the effect of coloration on out-strength.

The first panel of Fig. 6 shows that, except for the newly proposed `STRAND` measurement error model, all models fail to correctly detect the effect of coloration on out-strength. This failure is indicated by most methods having peak *p*values at values of *κ* > 0. The second panel of Fig. 6 shows that all models other than the `STRAND` measurement error model fail to correctly detect the effect of coloration on instrength. The third panel of Fig. 6 shows that all methods perform well across most of the parameter range: shyness has no effect on out-strength, and most methods return large *p*-values. However, for *κ* > 2, it appears that many methods begin to fail, suggesting that higher levels of censoring may significantly impact these results. The fourth panel of Fig. 6 shows that all models correctly indicate no effect of shyness on in-strength.

**Figure 6.**
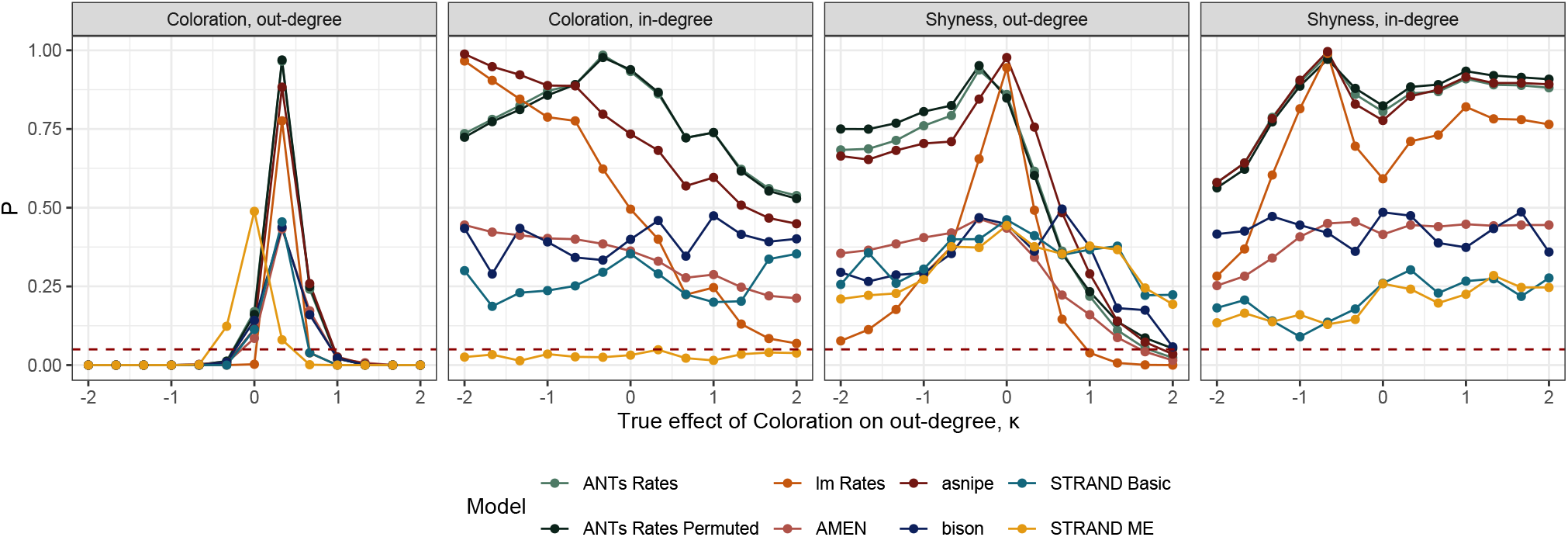
Analysis of simulated datasets subject to censoring, using eight different network approaches. See the caption to Fig. 4 for a description of how to interpret each panel below. In terms of performance, only the STRAND model that accounts for measurement error permits accurate estimation of the effect of coloration on out-strength in panel one. All other methods have peak *p*-values at values of κ > 0 and fail to detect true effects in the range of 0.5 < κ < 1.0. In panel two, only the STRAND model which accounts for measurement error accurately detects the true effect of coloration on in-strength. In panel three, all methods perform well across most of the parameter range, but only the STRAND models continue to suggest there is no effect of shyness on out-strength for values of κ > 1.5. In panel four, all models correctly indicate that there in no effect of shyness on in-strength.

### Robustness checks

Although the findings in Figs. 4–6 are fairly general, the relative performance of different modeling approaches might depend on the topology of the particular networks we used for the examples. In order to investigate the robustness of our main results, we simulate 200 random network data sets (each comprised of *n* = 65 nodes, and thus 2,080 dyads) for each of four levels of sampling bias and censoring intensity (i.e., 200× 4× 2 = 1, 600 data-sets in total). Within each condition, the networks are generated assuming: (1) a constant effect of coloration on out-strength of 1.75 (a strong effect), (2) a constant effect of coloration on instrength of 0.75 (a weak effect), (3) no effect of shyness on out-strength, and (4) no effect of shyness on in-strength. All other parameters in the model—including within- and between-block tie probabilities, the correlation between coloration and shyness, the effect a dyadic third variable, the strength of generalized and dyadic reciprocity, and the variance in sender, recipient, and dyadic random effects— are sampled randomly over wide, but empirically plausible ranges. Each of the eight analysis protocols are fit to each simulated network dataset and effects of coloration and shyness on in-strength and out-strength are estimated. We follow the commonly-used, but arbitrary decision rule that ‘accepts’ effects with *p* < 0.05 and considers effects with *p* > 0.05 to be ‘insignificant’. Then, we evaluate whether or not each method correctly recoveres the true positive effects of coloration on out-strength and in-strength, and correctly inferres there is no evidence for significant effects of shyness on out-strength or in-strength. Figs. 7 and 8 plot the results for sampling bias and censoring conditions, respectively. The Bayesian methods, STRAND and Bison, clearly outperform other methods in recovering small, true effects. The eight methods otherwise perform similarly: all methods recover large, true effects at similar rates and conclude there is no evidence for null, true effects at similar rates.

**Figure 7.**
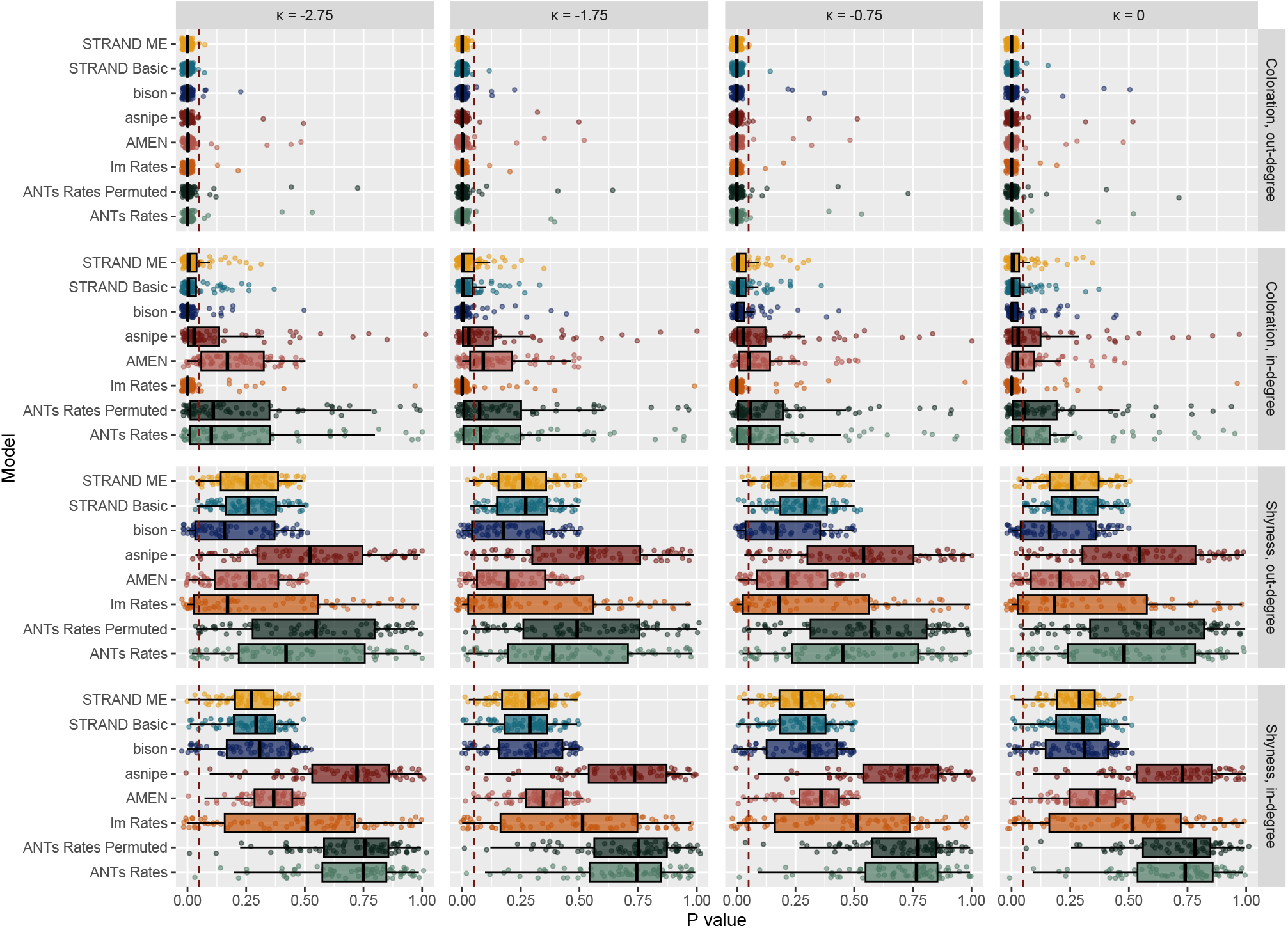
Robustness analysis using eight different network approaches to analyze simulated network data subject to biases in sampling intensity. In each of four conditions (where the effect of coloration on sampling intensity is set to: 0.00, -0.75, -1,75, and -2.75, respectively), 200 networks are generated assuming: (1) a constant effect of coloration on out-strength of 1.75 (a strong effect), (2) a constant effect of coloration on in-strength of 0.75 (a weak effect), (3) no effect of shyness on out-strength, and (4) no effect of shyness on in-strength. All other parameters in the generative model are randomized, so as to generate networks with wildly different topologies. Well-performing models should have all points (and boxplots) to the left of the vertical dashed line at *p* = 0.05 in the upper two panels, and to the right of that same line in the bottom two panels. In terms of correctly identifying the true effect of coloration on out-strength (a strong effect), most methods perform reasonably well, even with high levels of sampling bias. In terms of correctly identifying the true effect of coloration on in-strength (a weak effect), the Bayesian methods perform well, even with high levels of sampling bias. Permutation methods perform poorly, even when sampling bias is zero, and their performance only declines as sampling bias increases. In the bottom two panels, most methods correctly indicate that there is no reliable effect of shyness on either out-strength or in-strength.

**Figure 8.**
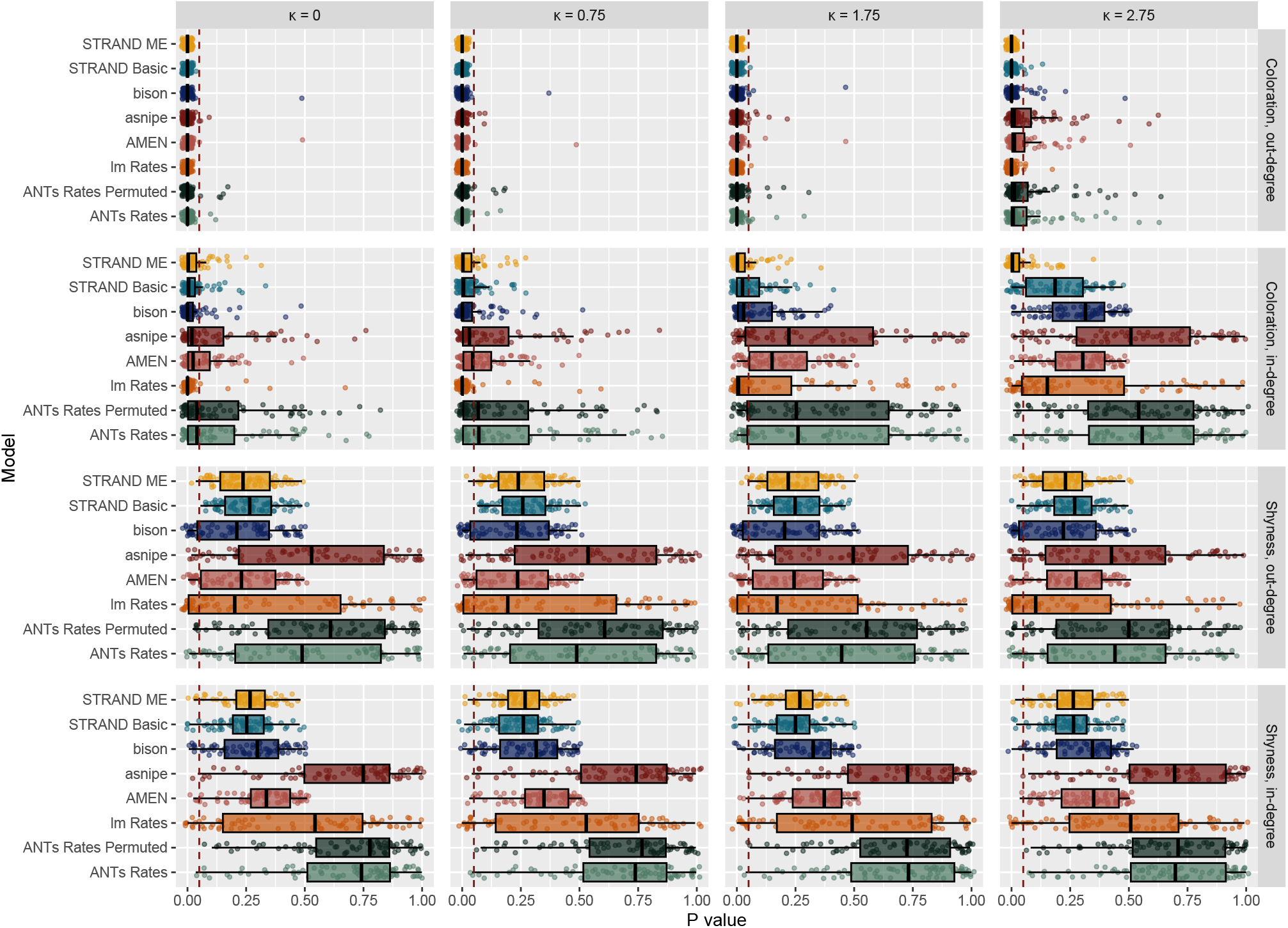
Robustness analysis using eight different network approaches to analyze simulated network data subject to biases in censoring. In each of four conditions (where the effect of coloration on censoring intensity is set to: 0.00, -0.75, -1,75, and -2.75, respectively), 200 networks are generated assuming: (1) a constant effect of coloration on out-strength of 1.75 (a strong effect), (2) a constant effect of coloration on in-strength of 0.75 (a weak effect), (3) no effect of shyness on out-strength, and (4) no effect of shyness on in-strength. All other parameters in the generative model are randomized, in order to generate networks with wildly different topologies. Well performing models should have all points (and boxplots) to the left of the vertical dashed line at *p* = 0.05 in the upper two panels, and to the right of that same line in the bottom two panels. In terms of correctly identifying the true effect of coloration on out-strength (a strong effect), most methods perform well in the absence of censoring issues. As the strength of censoring increases sufficiently, the permutation methods begin to fail at appreciable rates, while the Bayesian methods persist in yielding correct inference. In terms of correctly identifying the true effect of coloration on in-strength (a weak effect), the permutation methods perform poorly, even when censoring is zero, and their performance only declines as censoring increases. For small levels of censoring, all Bayesian methods yield correct inference. However, for high levels of censoring, only the STRAND model which accounts for measurement error permits accurate inference. In the bottom two panels, most methods correctly indicate there is no reliable effect of shyness on either out-strength or in-strength.

To visualize the frequency of inference errors, we process the data from Figs. 7 and 8 and plot the fraction of datasets for which application of a given analysis method led to incorrect inference in Fig. 9. An error rate of 0.05 is visualized as a dashed vertical line. In terms of detecting strong, true effects of coloration on out-strength, all methods have error-rates of less than 0.05 under most contexts.

**Figure 9.**
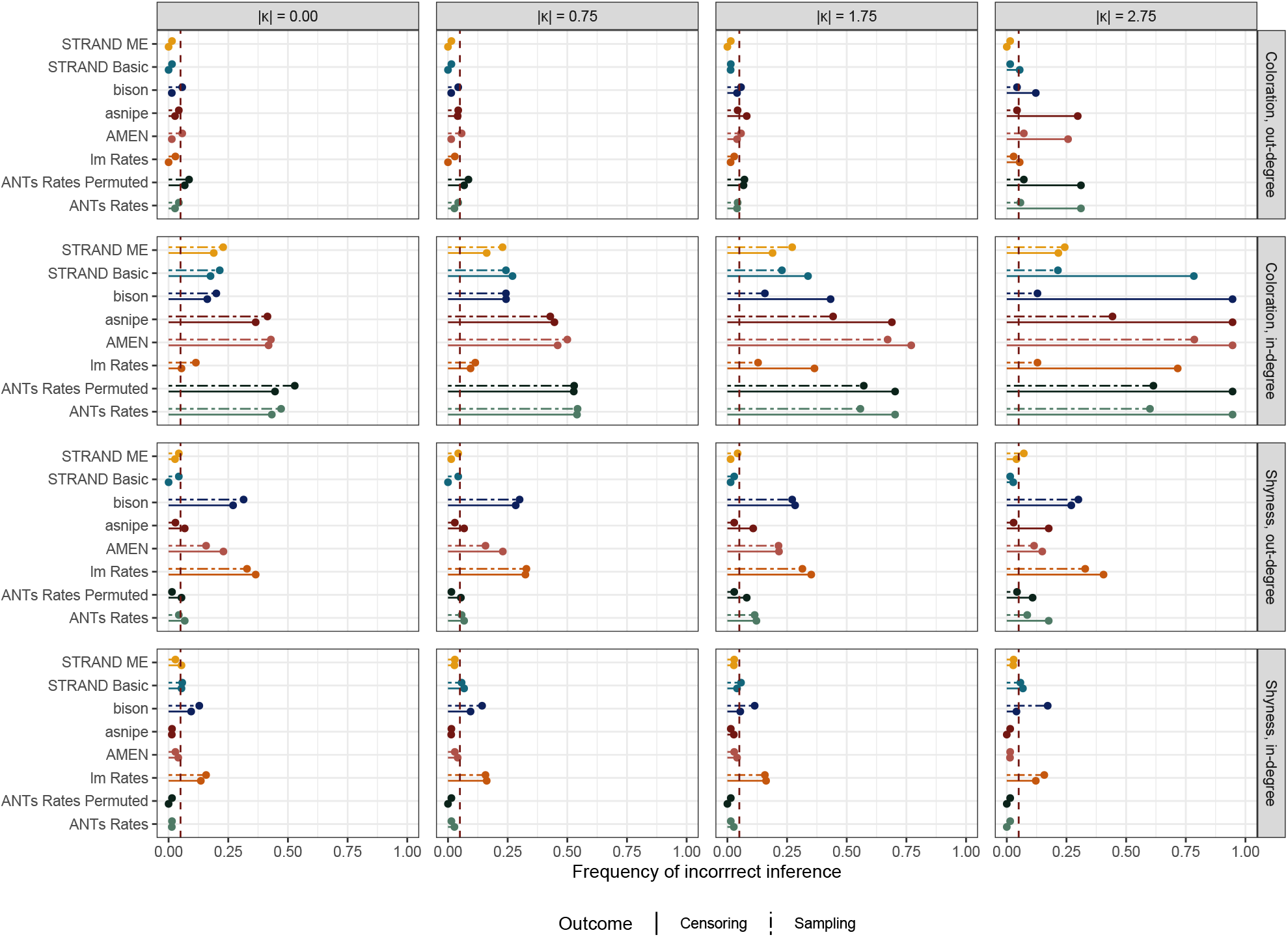
Robustness analysis using nine different network approaches with a simulated network of *n* = 65 nodes. In each of four conditions (where the effect of coloration on censoring probability is set to: 0.00, 0.75, 1,75, and 2.75, respectively), 200 networks are generated assuming: (1) a constant effect of coloration on out-degree of 1.75 (a strong effect), (2) no effect of shyness on out-degree, (3) a constant effect of coloration on in-degree of 0.75 (a weak effect), and (4) no effect of shyness on in-degree. All other parameters in the generative model are randomized to generate networks with wildly different topologies. In general, most methods are effective in not falsely inferring effects of shyness when none exist in the generative model. However, in the presence of high levels of censoring, the STRAND models have lower error rates than the other methods. In terms of correctly identifying the true effect of coloration on out-degree (a strong effect), most methods perform reasonably well in the absence of censoring issues. Within each condition the permutation methods have the highest error rates and the STRAND models the lowest. As the intensity of censoring increases, all methods other than STRAND fail to detect strong, true effects of coloration on out-degree nearly a third of the time. Lastly, all models struggle to identify the true effect of coloration on in-degree, as this effect was purposely set to be weak and hard to detect given the sample size of *n* = 65. Across conditions, however, the STRAND models perform appreciably better than all other methods.

Only the permutation methods in high censoring contexts have error rates appreciably higher than 0.05. In contrast, in terms of correctly inferring no reliable effect of shyness on out-strength and in-strength, the lm and Bison packages had abnormally high error rates. The reason for this is that these two approaches do not incorporate the random-effects structure of the Social Relations Model, thus they overestimate the weight of evidence in favor of small effects. The `STRAND` models, in contrast, are well titrated: the error rates are generally less than 0.05, except in the case of a weak effect of coloration on in-strength in the presence of strong data censoring, where the error rates are closer to 0.25. This case, however, was intended to be difficult for all methods. Moreover, the `STRAND` models performed better here than all other methods, especially when sampling bias and censoring effects were present.

### Empirical applications

Lastly, we highlight the value of our model for inference using an empirical dataset. We draw on data published by Gelardi et al. (2020), who simultaneously measured Baboon social networks using both direct observation and RFID collars. In this dataset, the direct observation data are more nuanced, in that they represent directed behaviors between individuals *i* and *j*, and distinguish types of social relationships (e.g., grooming, social resting, and playing). The direct observation data, however, are subject to censoring: 18% of focal observations with duration information are labeled as ‘focal not visible’. An important fraction of social interactions might thus occur in locations where the focal could not be observed by the researcher. Moreover, some individuals tend to be unobservable at higher rates than others. Data from the RFID collars, in contrast, represent simple proximity events and can only be used to construct undirected contact networks. The benefit of the RFID data, however, is that they are robust to censoring, in that contact information is recorded consistently across individuals, even if contact events happen in locations where direct observations could not be made.

To apply our model, we first subset the Gelardi et al. (2020) data to include time points where both direct observation and RFID tracking were performed simultaneously. We then calculate individual differences in censoring probability by evaluating the relative counts of interactions involving each individual in the observational and RFID datasets. We assume that RFID detection events give unbiased rates of inter-individual contact, and therefore that differences in contact rates between the two networks are driven by interindividual differences in observability in the observational study. For example, if individuals *i* and *j* are observed 850 and 640 times in the RFID data, and 330 and 175 times in the observational data, then we can conclude that *i* and *j* have different observability rates of 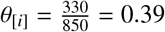 and 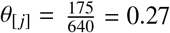 . However, since the RFID and direct observation methods differ in terms of sampling rate, *θ* is only indicative of *relative* observability rates between individuals. As such, we consider two arbitrary normalizations of *θ*, one scales *θ* such that its maximal element equals 0.6, and the other scales *θ* such that its maximal element equals 0.99. The censoring rate vector is then *η* = 1 − *θ*.

For this example, we recode the observational data to be undirected—by adding the outcome and exposure matrices to their own transposes—so that we can directly compare adjacency matrices between the observational and RFID studies. Finally, we estimate the weighted adjacency matrices describing latent between-individual tie strengths using: (1) the RFID data alone, (2) the observational data alone, and (3) the observational data combined with censoring rate estimates. We then evaluate the accuracy of our models by comparing the observational networks—with and without censoring adjustment—to the RFID ground-truth rates.

In Figure 10, we plot the results. In frame 10a, we see the adjacency matrix implied by the RFID data alone. In frame 10b, we see the adjacency matrix implied by the observational data alone. We note that the observational adjacency matrix appears too sparse; that is, there are true contact events—often corresponding to weaker strength ties—in the RFID data that are not detected as readily via focal observation. In frames 10c–10d, we plot the adjacency matrices implied by the observational data in combination with data on censoring rate. Here we note a much closer correspondence between the observational adjacency matrices and the ground-truth RFID adjacency matrix.

**Figure 10.**
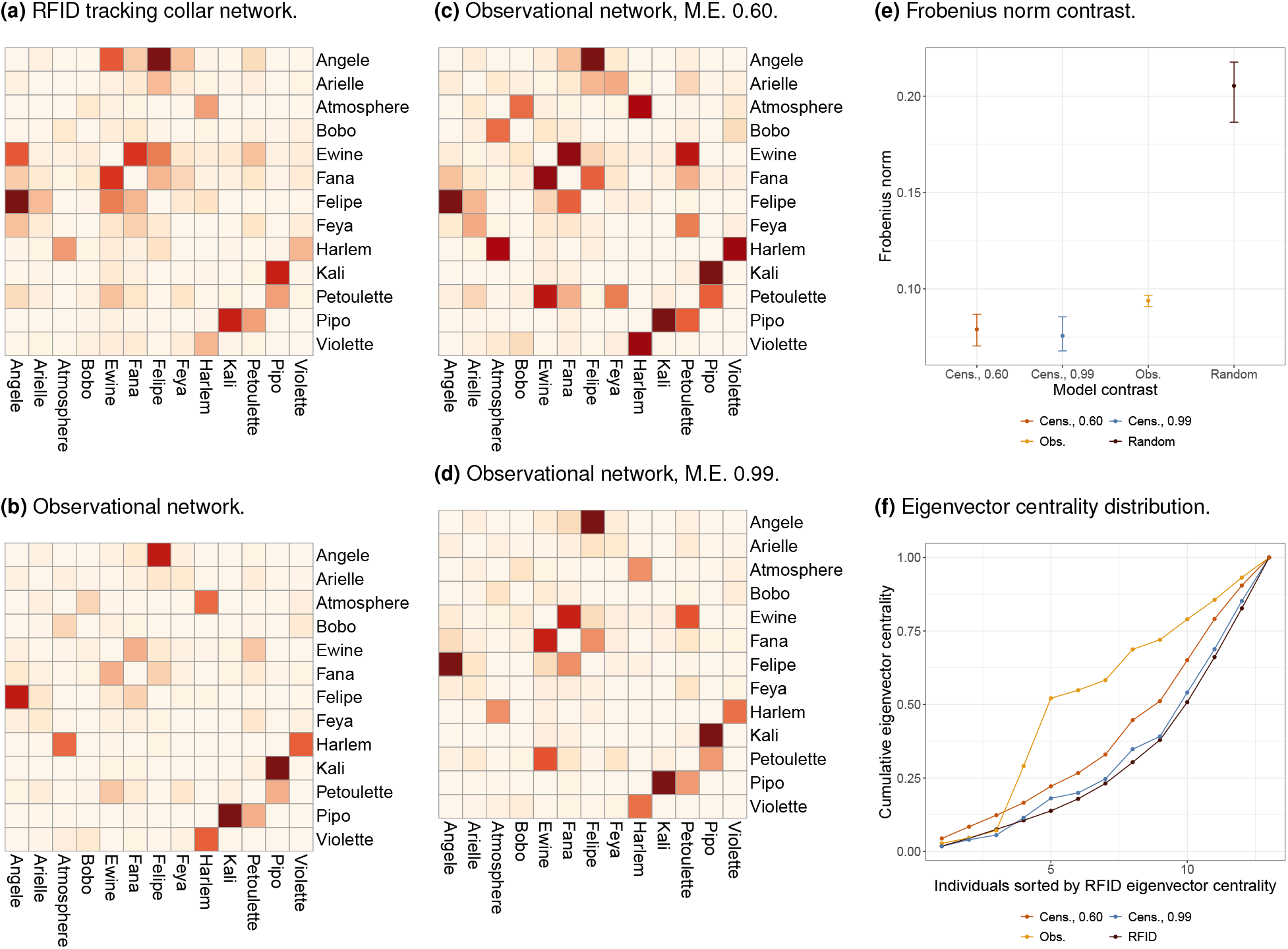
Network estimation using data from Gelardi et al. (2020). Frames 10a–10d plot the adjacency matrices recovered under different methods. Darker red indicates higher strength ties between individuals. Frame 10a is based on RFID data alone, frame 10b is based on observational data alone, and frames 10c and 10d are based on observational data with adjustment for measurement error. Frame 10e plots the Frobenius norm between the RFID adjacency matrix in frame 10a and the other three adjacency matrices in frames 10b–10d. A smaller Frobenius norm indicates more similar matrices. Lastly, 10f plots the normalized cumulative eigenvector centrality of individuals, sorted in order of increasing eigenvector centrality in the RFID data. By accounting for measurement error in the observational network, we are better able to recover the eigenvector centrality distribution apparent in the unbiased RFID network.

To formally measure the improvement, we calculate the Frobenius norm (see: Cui et al. 2016) between the RFID adjacency matrix and each of the other matrices, including a random permutation of the observational adjacency matrix (see Figure 10e). We note first that the Frobenius norm between the RFID adjacency matrix and each of the observational adjacency matrices is quite small in comparison to the random matrix. This shows that observational and RFID networks are more similar in structure that expected by chance. We also see a moderately sized, but highly reliable, reduction in the size of the Frobenius norm upon application of the measurement error model. This indicates that control for censoring improves identification of network structure.

More broadly, non-linear, network-wide properties, such as the eigenvector centrality distribution, are widely known to be sensitive to even small changes in network topology (Cavallaro et al. 2024). As such, even moderately sized improvements in the estimation of latent network structure can have substantial impact on recovery of higher level network properties. In Figure 10f, we plot the normalized cumulative eigenvector centrality of individuals, sorted in order of increasing eigenvector centrality in the RFID data. We note a substantial divergence in the eigenvector centrality distribution between the RFID network and the observational network. However, by accounting for censoring, we are able to almost perfectly recover the RFID eigenvector centrality distribution using only the observational data and estimates of censoring rate.

## Discussion

Establishing the reliability of methodologies in Animal Social Network Analysis (ASNA) is crucial for their effective application in empirical studies of social behavior. In this study, we developed a novel Bayesian generative model that estimates network structure (STRAND ME) and explicitly estimates and adjusts for censoring biases. Additionally, we developed a simulation approach to systematically evaluate our model and compare it to multiple ASNA methods under various types and intensities of measurement biases. Unlike the previous simulation studies discussed in the introduction, we simulated sampling and observational biases independently by generating probabilities for over- or under-sampling (to generate sampling bias) and/or censoring (to generate observation biases). In doing this, our objective was to understand the applicability and limitations of ASNA methods, while also refining and optimizing these methodologies to enhance their reliability for real world studies of animal social networks.

The model that we have developed here estimates and adjusts for the probability of detectability for each individual. Through this approach, our model assumes that social ties may not be observed between two individuals if either one of the two individuals is undetectable during an observation—even if a social tie does truly exists. The model formalises this by introducing a parameter that represents the probability of detecting an individual, which may vary based on individual-specific characteristics (e.g., more cryptic colors in females birds: Farine and Whitehead 2015a). As a result, our approach recovers the true underlying social connections and effectively corrects for biases introduced by imperfect observation. Because of this, our model outperforms all other ASNA methods included in the study, as demonstrated in Scenario 3. Moreover, our model’s flexibility allows it to be coupled with models of network structure, such as the social relations model (Kenny and La Voie 1984), stochastic blockmodel (Holland et al. 1983), or a combination of the two (Redhead et al. 2023b). This flexibility in model specification is a key strength, as it enables the integration of individual-level random effects, dyad-level random effects, and covariates at both of these levels to capture theoretically important features of the network. By incorporating these parameters, our model estimates important structural features, while ensuring that measurement biases common to ASNA are accounted for.

By comparing several popular ASNA methods, our simulation experiments highlight the utility of recently proposed Bayesian approaches for understanding animal social relationships. These Bayesian models perform particularly well in addressing the data collection biases that are common to empirical studies of animal sociality. Specifically, our results show that STRAND and BISON are capable of handling sampling biases, demonstrating improved performance compared to permutation based approaches (i.e., ANTS rates, ANTS rates permuted, lm rates, and MRQAP) in nearly all scenarios. The improved performance of these approaches is likely due to two major strengths of Bayesian generative models of network structure. First, as outlined in the introduction, Bayesian methods improve statistical reasoning by scaling the influence of each data point according to the sampling effort, unlike traditional indices of sociality. This adjustment allows for a more accurate representation of the data, particularly in cases with uneven sampling. Second, Bayesian methods offer greater flexibility than permutation approaches. For example, Bayesian generative models of network structure such as STRAND, allow for the integration of individual-level parameters that are analogous to in-strength and out-strength—as well as dyad-level parameters—within the same regression model. This capability leads to more accurate results when testing the effects of individual characteristics on multiple network measures. In addition, it addresses potential biases that traditional methods might overlook.

In the presence of sampling biases, simple linear regression techniques—with or without permutations—begin to falter when evaluating the effects of individual characteristics on multiple network measures. As demonstrated in Scenarios 1 and 2, these approaches can accurately estimate the effect of coloration on out-degree. However, they tend to overestimate the effect on in-strength once the *κ* value reaches -1 (i.e., when there is a large, true effect of colouration on out-strength). This overestimation arises because these approaches run separate linear regression models for in-strength and out-strength. Such an approach fails to account for the potential dependencies between these two variables. More specifically, when one variable is omitted from the model, the regression may incorrectly attribute variation in the dependent variable to the independent variable without considering any shared variance with the omitted variable. Indeed, we observe that as the relationship between coloration and out-strength increases, the regression model interprets the shared variance between in-strength and out-strength as evidence of a direct effect of coloration on in-strength. Consequently, the regression coefficients associated with coloration could become inflated, leading to an overestimation of its influence on in-strength. To mitigate the risk of misattribution and omitted variable bias, an approach that aims to appropriately capture the structural features of a network is necessary (Snijders 2011). This could be, for example, a model that estimates the probability of social ties—which includes parameters estimating both in-strength and out-strength, alongside other theoretically relevant features of a network—as done in Bayesian generative models of network structure (e.g., STRAND).

However, it is important to note that all statistical models, including Bayesian ones, require careful attention to the details, in order to ensure that a model accurately reflects the generative processes underlying the data. Indeed, we tested 3 different Bayesian generative models of network structure with different generative processes and our results highlight certain limitations of the BISON and AMEN approaches within our simulation scenarios. Specifically, at high values of *κ* (i.e., when there is a large, true effect of *coloration* on out-strength), the BISON approach can produce false positive estimates for the relationship between *shyness* and out-strength across Scenarios 1, 2, and 3. Similarly, the AMEN approach also shows false positives for the relationship between shyness and out-strength in all three scenarios, and it additionally suffers from false positives in estimating the relationship between coloration and out-strength. Our robustness analysis emphasizes the importance of incorporating structural parameters that appropriately represent the data (e.g., the social relations model parameters in STRAND). Without these structural parameters, models like BISON tend to overestimate the weight of evidence in favor of small effects. These findings suggest that only our new STRAND model, which accounts for both sampling and censoring biases, accurately estimates the effects of the various factors in the presense of data censoring.

However, the improved performance of the STRAND models in cases without data censoring are partially due to the fact that our simulations featured a data generating procedure that STRAND was specifically developed to account for—i.e., a case where there is correlated node-level heterogeneity in degree, and dyad-level correlations in tie strength. We view these features as near omnipresent in emprical data-sets, however, and so we do not see our simulation as artifically ‘stacking the deck’ in favor of methods based on the social relations model. Nevertheless, we note that although the BisonR tutorial^§^ for dyadic regression we followed did not include both dyadic and node-level random effects, such models should eventually be possible to code within the Bison framework.

Here, we propose a robust approach to animal social network anlysis that effectively handles sampling and censoring biases that are well-described threats-to-infernce in real-world research projects; our model is thus tailored to these specific issues. Consequently, researchers may need to further develop and test bespoke statistical models to address other additional biases—a task that is currently only implementable when using an analysis framework based on non-null models of the data-generating process.

Overall, our research highlights that Bayesian generative models of network structure are particularly well-suited approaches for ASNA. Indeed, Bayesian methods offer greater flexibility than permutation approaches, e.g., by permitting the joint assessment of in-strength and out-strength within the same model, and by permitting the simultaneous estimation of network features and interaction strengths. Additionally, Bayesian generative models of network structure enable the incorporation of additional effects—such as stochastic blockmodels for group-structure—and offer the flexibility to integrate prior knowledge and uncertainty into the analysis. This integrated approach leads to more robust models.

Finally, despite the numerous advantages of Bayesian modeling, practical implementation can pose challenges. For instance, while we have successfully incorporated censoring control within a Bayesian framework, its application remains complex due to the need to estimate a censoring effect—which requires additional data-sources or experiments. One potential approach, as we showed in the empirical example, is to evaluate the dissimilarity between observations from human observers and biologgers (e.g., RFID or GPS trackers; assuming that biologging is unbiased with respect to individuals). However, this approach may need to be conducted at each observation period, such as annually in a longitudinal study, since group composition can change. The dependence on additional data may significantly complicate the practical implementation of our measurement-error model; however, our simulation study shows that without accounting for censoring when it is present, valid inference might not be possible . Therefore, alternative strategies for assessing data censoring might be needed. For example, comparing scan samples with focal-follow samples, where focal samples may be less biased by censoring (Gilby et al. 2010), could be a viable solution.

Another challenge with Bayesian frameworks, particularly for empirical researchers who may not have advanced training in statistical programming, is the current state of Bayesian analysis software. Current Bayesian frameworks, such as those built using Stan (Stan Development Team 2021), are highly specialized and require statistical expertise that is not typically part of the training for many researchers. While the flexibility of Bayesian approaches allows for the development of custom models tailored to specific research questions, the complexity of these models—coupled with the lack of training in statistics and programming for empirical researchers—makes their use difficult. Future developments in Bayesian analysis software should focus on expanding the range of effects and parameter specifications that can be flexibly incorporated into out-of-the-box models (e.g., in a similar vein to the RSiena software: Ripley et al. 2011) and STRAND, Bison, and AMEN. This would enable researchers to readily integrate components into custom models, providing greater control and adaptability in model specification. Addressing these practical challenges and advancing Bayesian frameworks could greatly advance ASNA research.

In conclusion, our study here offers some insight into the reliability and limitations of ASNA methodologies across various types and intensities of data-collection biase. Our results underscore the need for the field of ASNA to progress beyond regression analyses and permutation methods and apply generative models of network structure that are tailored to specific research questions, as well as to any measurement biases that characterise a given dataset. Here we have introduced a Bayesian generative model of network structure that effectively addresses both sampling and observational biases while estimating network structure. By systematically evaluating this model and comparing it to existing methods, we provide a more robust and accurate method for analysing animal social networks.

## Data Availability

The code used to reproduce the analysis and manuscript is available at: https://github.com/BGN-for-ASNA/ASNA_reliability.

## Author Contributions

Sebastian Sosa conceived the research idea, developed the code, and coauthored the manuscript. Mary Brooke McElreath contributed to the manuscript writing. Daniel Redhead reviewed the code and contributed to the manuscript writing. Cody T. Ross developed the code and coauthored the manuscript.

## Footnotes

§ https://jhart96.github.io/bisonR/articles/getting_started.html

* For proof, consider observing *x*_[i,j]_ directed outcomes, given *E*_[i,j]_ opportunities to observe those outcomes. Then assuming flat prior information over the unit interval, the posterior probability of directed outcomes, Π_[i,j]_, is given by a Beta distribution (Perks 1947), with probability density: 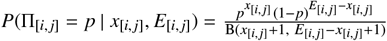 . Note that this expression is not equivalent to the index value of 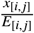 for any finite data-set.

† To apply the model we describe, one must round any continuous measures to integer units.

‡ For standard estimates, we report the standard *p*-values. For methods involving permutations, we report the permutation *p*-values. For Bayesian estimates, we report the probability of a sign error, which is defined as: 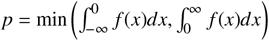, where *f* (*x*) is the posterior probability density that the coefficient *β* = *x*.

## Notes

https://github.com/BGN-for-ASNA/ASNA_reliability

### Competing Interest Statement

The authors have declared no competing interest.

